# The crosstalk between the anterior hypothalamus and the locus coeruleus during wakefulness is associated with low frequency oscillations power during sleep

**DOI:** 10.1101/2025.04.15.647930

**Authors:** Nasrin Mortazavi, Puneet Talwar, Ekaterina Koshmanova, Roya Sharifpour, Elise Beckers, Ilenia Paparella, Fermin Balda, Christine Bastin, Fabienne Collette, Laurent Lamalle, Christophe Phillips, Mikhail Zubkov, Gilles Vandewalle

## Abstract

**Study Objectives:** Animal research has demonstrated that sleep regulation heavily depends on a network of subcortical nuclei. In particular, whether the crosstalk between the Locus Coeruleus (LC) and hypothalamic nuclei influences sleep variability and age-related changes in humans remains unexplored. This study investigated whether the effective connectivity between the LC and subparts of the hypothalamus is associated with the electrophysiology of rapid eye movement sleep (REMS).

**Methods:** Thirty-three healthy younger (∼22y, 27 women) and 18 older (∼61y, 14 women) individuals underwent 7-Tesla functional magnetic resonance imaging during wakefulness to investigate the effective connectivity between LC and distinct hypothalamus subparts encompassing several nuclei. Additionally, we recorded their sleep electroencephalogram (EEG) to explore relationships between effective connectivity measures and REMS theta energy and sigma power prior to REMS episodes.

**Results:** The effective connectivity analysis revealed robust evidence of a mutual positive influence between the LC and the anterior-superior and posterior hypothalamus, supporting the idea that the connectivity patterns observed in animal models are also present in humans. Furthermore, our results suggest that in older adults, stronger effective connectivity from the anterior-superior hypothalamus, including the preoptic area, to the LC is associated with reduced REM theta energy. Specificity analysis showed that this association was not limited to REM theta energy but also extended to specific lower-frequency bands during REMS and NREMS.

**Conclusions:** These findings highlight the complex age-dependent modulation of the LC circuitry and its role in sleep regulation. Understanding these neural interactions offers valuable insight into the mechanisms driving age-related sleep changes.

## Introduction

Sleep and wakefulness regulation and the fine-tuning of vigilance state are regulated by a circuit of subcortical nuclei located in the basal forebrain, thalamus, hypothalamus and brainstem as part of the ascending activating system ^1–3^. The locus coeruleus (LC), in the brainstem, consists of the main source of norepinephrine (NE) of the brain and sends widespread monosynaptic projections to nearly all brain regions ^4,5^. Research in animal models showed that its activity must decrease for transition from wakefulness to sleep. During sleep, the LC shapes the switch between slow wave sleep (SWS) and rapid eye movement sleep (REMS) as well as some of the microstructure elements of sleep ^6^. Investigation in humans indicated that degeneration of the LC is likely driving part of the alteration in sleep-wake regulation commonly found in healthy and pathological aging^7^. Likewise, inappropriate LC activity during both wakefulness and sleep is also probably key to the emergence and maintenance of insomnia ^8^. LC hyperactivity would contribute to a state of hyper arousal during wakefulness while it would either reduce REM sleep occurrence or REM bout stability during sleep, two phenomena associated with insomnia disorder ^9^. In line with this hypothesis, we recently reported that a balanced activity of the LC during wakefulness is associated with a more intense REMS, as indexed by the overnight energy over REM most typical oscillatory mode (theta oscillations) ^10,11^. We further showed that LC activity during wakefulness was associated with the sigma power immediately preceding REM sleep episode ^11^, a sleep feature causally linked with the LC in animal models ^12^.

Research in animal models also established that several nuclei of the hypothalamus are key elements of the circuit regulating sleep and wakefulness. The posterior part of the hypothalamus includes the lateral hypothalamus (LH) and the tuberomammillary nucleus (TMN) which produce orexin and histamine, respectively, two neuromodulators stimulating wakefulness, while the LH further produces melanin-concentrating hormone (MCH) which is considered to stabilize sleep and REM sleep in particular ^3^. The lateral hypothalamus is further known to exert an excitatory influence over the LC through glutamatergic afferents ^13^. The superior part of the anterior hypothalamus includes preoptic nuclei inhibiting the nuclei of the ascending activating system, including the LC, notably through the production of gamma-aminobutyric acid (GABA) and favor sleep initiation ^2^. The inferior part of the anterior hypothalamus further includes the main circadian clock – in the suprachiasmatic nucleus - which organizes sleep and wakefulness in time of the 24h light-dark cycle mostly through vasoactive intestinal peptide (VIP) and GABA ^14^, and notably through indirect projection to the LC ^15^. Similar to the LC, the nuclei of the hypothalamus undergo age-related changes, which can disrupt their regulatory role in sleep, potentially contributing to altered sleep patterns in healthy and pathological aging ^16^.

As for the LC, most of our understanding of the roles of the hypothalamus nuclei in sleep and wakefulness regulation arises from animal studies such that translations to human beings are needed if one wants to develop efficient interventions geared toward the neuromodulator systems underlying vigilance state in healthy individuals and patients. Whether the crosstalk between the LC and nuclei of the hypothalamus may underly variability in sleep electrophysiology and changes in aging has not been investigated.

Similar to our initial reports of an association between LC activity and REM sleep, we hypothesized that the cross-talk between the LC and nuclei of the hypothalamus during wakefulness would reflect the overall variability of this cross-talk, including during sleep ^10,11^. We used the same 7 Tesla functional magnetic resonance imaging (7T fMRI) dataset as our initial studies ^10,11^ to test whether effective connectivity between the LC and subparts of the hypothalamus would be related to REM theta energy and sigma power prior to REM episodes, i.e. the sleep parameters related to wakefulness LC activity in our previous reports ^10,11^. We anticipated that the connectivity between the LC and at least the anterior-superior, anterior-inferior or the posterior-lateral subpart of the hypothalamus would be markers of REM sleep quality. We further anticipated that the associations would be affected by aging.

## Methods

This study was approved by the faculty-hospital ethics committee of ULiège. All participants provided written informed consent and received financial compensation. The study is part of a larger project that has led to previous publications.^10,11,17^ Most of the methods were described in details in^10,11^.

### Participants

Fifty-two healthy participants were included in the study. Due to technical issues, one subject was excluded and the final sample included 51 participants, with 33 healthy young (22.2±3.2y, 27 women) and 18 late middle-aged (60.9±5.4y, 14 women) individuals **(Table 1)**. The exclusion criteria were as follows: history of major neurologic/psychiatric diseases or stroke; recent history of depression/anxiety; sleep disorders; medication affecting the central nervous system; smoking, excessive alcohol (>14 units/week) or caffeine (>5 cups/day) consumption; night shift work in the past 6 months; BMI ≤18 and ≥29 (for older individuals) and ≥25 (for younger individuals). All older participants had to show normal performance on the Mattis Dementia Rating Scale (score > 130/144) ^18^. Due to a miscalculation at screening, 1 older participant had a BMI of 30.9 and one of the younger participants had a BMI of 28.4. Since their data do not deviate substantially from the rest of the sample these participants were included in the analyses (including BMI as a covariate in our statistical models did not modify our results).

**Table 1.**
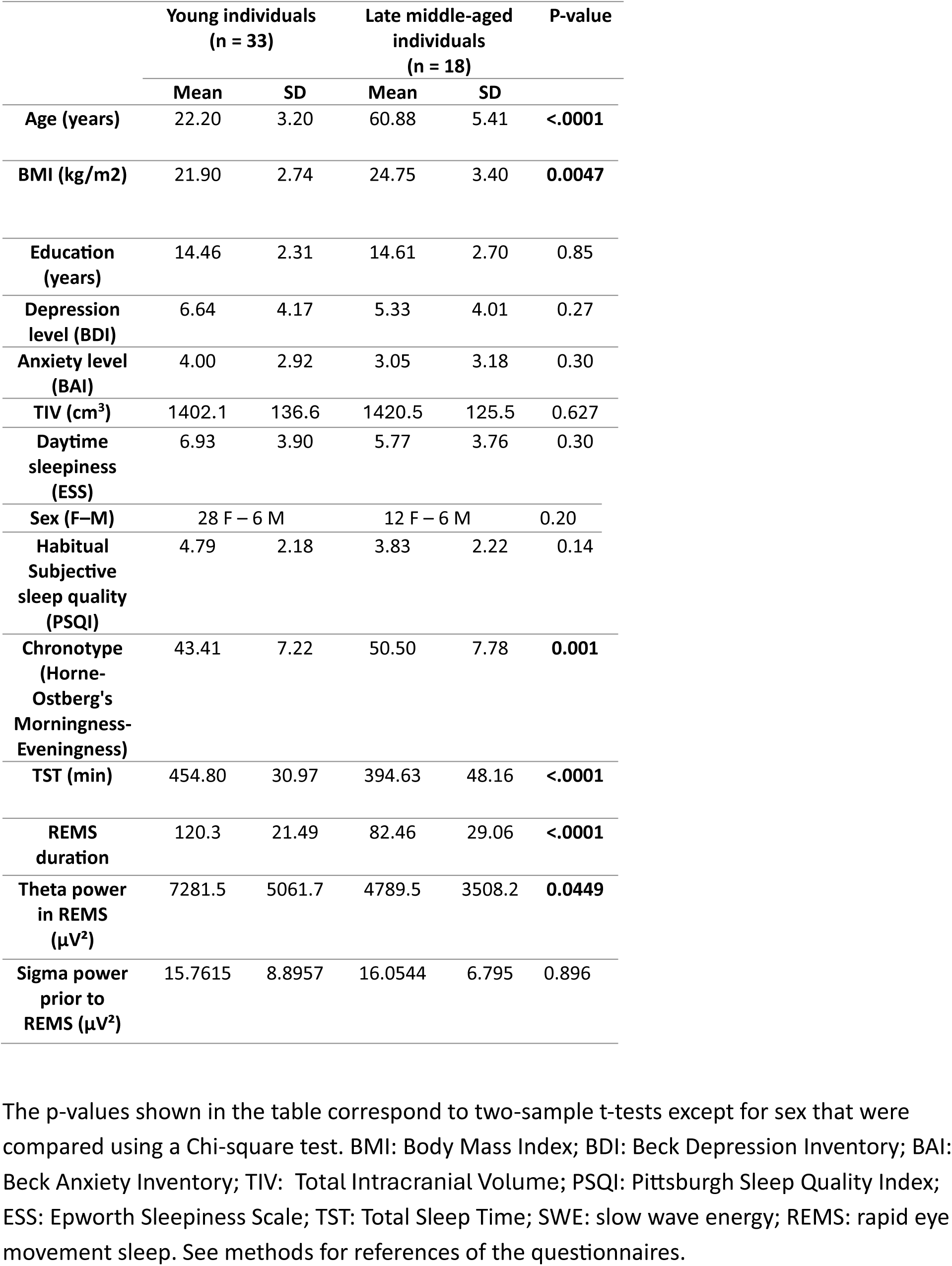
Characteristics of the study sample.

|  | Young individuals<br>(n = 33) |  | Late middle-aged<br>individuals<br>(n = 18) |  | P-value |
| --- | --- | --- | --- | --- | --- |
|  | Mean | SD | Mean | SD |  |
| Age (years) | 22.20 | 3.20 | 60.88 | 5.41 | **<.0001** |
| BMI (kg/m<sup>2</sup>) | 21.90 | 2.74 | 24.75 | 3.40 | **0.0047** |
| Education (years) | 14.46 | 2.31 | 14.61 | 2.70 | 0.85 |
| Depression level (BDI) | 6.64 | 4.17 | 5.33 | 4.01 | 0.27 |
| Anxiety level (BAI) | 4.00 | 2.92 | 3.05 | 3.18 | 0.30 |
| TIV (cm<sup>3</sup>) | 1402.1 | 136.6 | 1420.5 | 125.5 | 0.627 |
| Daytime sleepiness (ESS) | 6.93 | 3.90 | 5.77 | 3.76 | 0.30 |
| Sex (F–M) | 28 F – 6 M |  | 12 F – 6 M |  | 0.20 |
| Habitual Subjective sleep quality (PSQI) | 4.79 | 2.18 | 3.83 | 2.22 | 0.14 |
| Chronotype (Horne-Ostberg's Morningness-Eveningness) | 43.41 | 7.22 | 50.50 | 7.78 | **0.001** |
| TST (min) | 454.80 | 30.97 | 394.63 | 48.16 | **<.0001** |
| REMS duration | 120.3 | 21.49 | 82.46 | 29.06 | **<.0001** |
| Theta power in REMS (μV<sup>2</sup>) | 7281.5 | 5061.7 | 4789.5 | 3508.2 | **0.0449** |
| Sigma power prior to REMS (μV<sup>2</sup>) | 15.7615 | 8.8957 | 16.0544 | 6.795 | 0.896 |

The p-values shown in the table correspond to two-sample t-tests except for sex that were compared using a Chi-square test. BMI: Body Mass Index; BDI: Beck Depression Inventory; BAI: Beck Anxiety Inventory; TIV: Total Intracranial Volume; PSQI: Pittsburgh Sleep Quality Index; ESS: Epworth Sleepiness Scale; TST: Total Sleep Time; SWE: slow wave energy; REMS: rapid eye movement sleep. See methods for references of the questionnaires.

### Protocol

Participants’ sleep was recorded in the lab twice. During the first session, participants completed a night of sleep under polysomnography to screen for sleep abnormalities (apnea hourly index and periodic leg movement >15; parasomnia or REM behavioral disorder). All participants further completed a whole-brain structural MRI (sMRI) acquisition and a specific acquisition centered on the LC. Participants were then requested to sleep regularly for 7 days before the baseline night during the second session (±30min from their sleep schedule) based on their preferred schedule (compliance was verified using sleep diaries and wrist actigraphy - Actiwatch and AX3, AXIVITY LTD, Newcastle, UK). The evening before the baseline night, participants first completed questionnaires including Beck Depression Inventory (BDI)^19^, Beck Anxiety Inventory (BAI)^20^, the Pittsburgh Sleep Quality Index (PSQI)^21^, Epworth sleepiness scale (ESS)^22^ and Horne-Ostberg’s Morningness-Eveningness scale^23^ for assessing depression, anxiety, sleep quality, sleepiness, and chronotype respectively. They remained awake for 3h under dim light (<10 lux) for electrode placement and preparation to sleep prior to the recording of their habitual sleep in darkness under EEG. Approximately 3h after wake-up time under dim light (<10 lux), participants completed a functional MRI (fMRI) session that included 3 tasks (**Figure 1A**). This paper is centered on the analyses of the perceptual rivalry task and auditory salience detection task.

**Figure 1.**
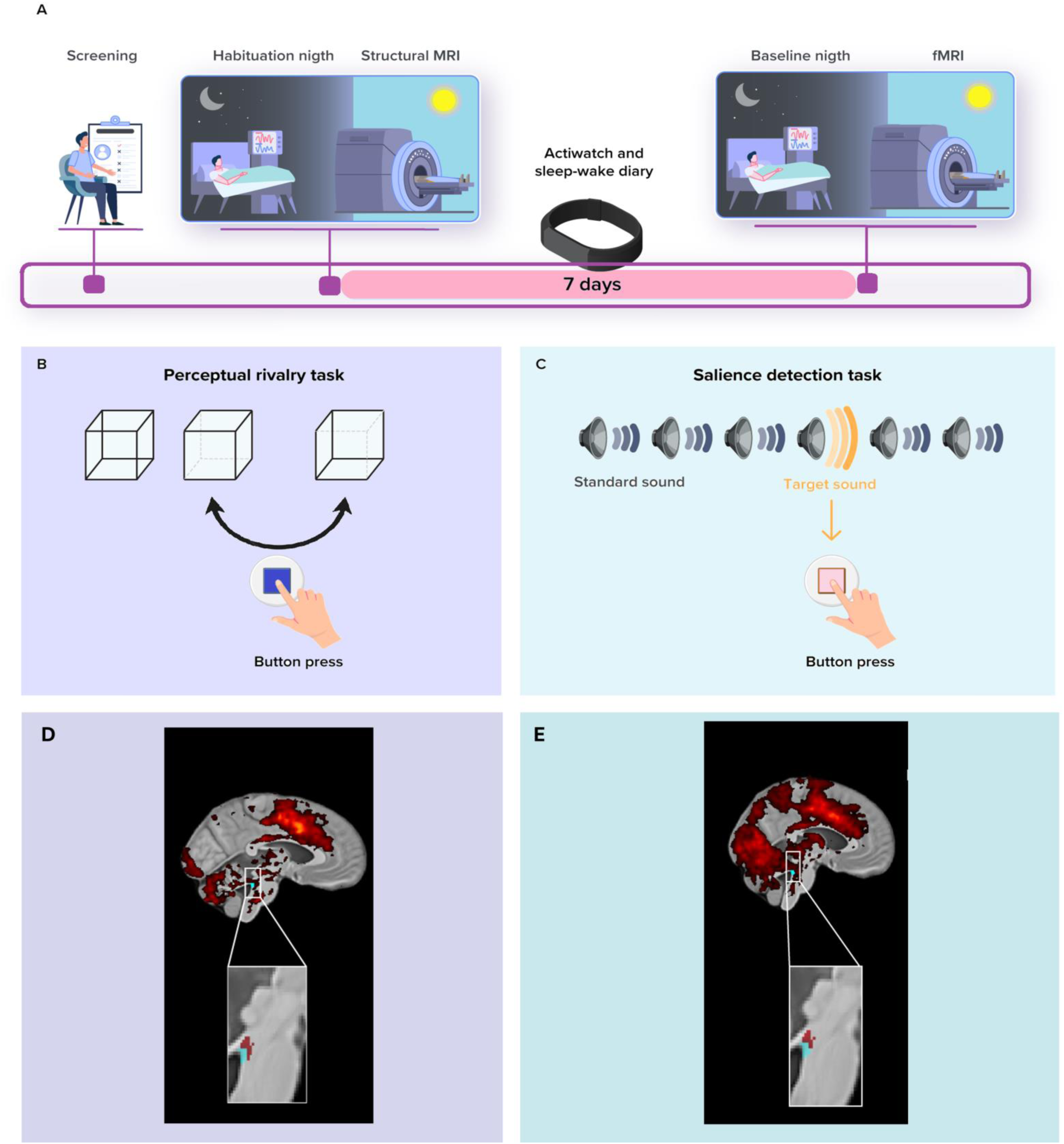
Overview of the study protocol. (A) After screening, participants completed an in-lab screening (i.e., habituation night) under polysomnography to minimize the effect of the novel environment for the subsequent baseline night and to exclude volunteers with sleep disorders. They further completed a structural 7T MRI session including a whole-brain structural MRI and a LC-specific sequence. After 7 nights of regular sleep-wake time at home, which was confirmed by actigraphy data and/or sleep-wake diary, participants came to the lab three hours before their sleep time and were maintained in dim light (<10 lux) until sleep time. Participants’ habitual baseline sleep was recorded overnight in-lab under EEG to extract our main sleep features of interest. All participants underwent an fMRI session approximately 3h after wake-up time (following ≥45min in dim light - < 10lux), during which they completed the visual perceptual rivalry task. (B) The Visual perceptual rivalry task consisted of watching a 3D Necker cube, which can be perceived in two different orientations (blue arrow), for 10 blocks of 1min separated by 10s of screen-center cross fixation (total duration ∼12min). Participants reported switches in perception through a button press. (C) The auditory salience detection task consisted of an oddball paradigm requiring button-press reports on the perception of rare deviant target tones (20% occurrence) within a stream of frequent tones (total duration ∼10min). (D) Whole-brain and LC responses to the perceptual switches during the visual perceptual rivalry task. [MNI coordinates: (−4 −37 -21 mm)]. The image at the top shows the whole-brain results using significance for a threshold of p< 0.05 FDR-corrected (t > 2.16) over the group average brain structural image coregistered to the MNI space. The inset at the bottom shows the LC probabilistic template (blue) created based on individual LC masks and the significant activation detected within this mask (red). The legend shows the t-values associated with color maps. (E) Whole-brain and LC responses to the target sound during the auditory salience detection task [MNI coordinates: (−4 −34 -21 mm)]. The image at the top show the whole-brain results using significance for a threshold of p< 0.05 FDR-corrected (t > 2.33) over the group average brain structural image coregistered to the MNI space. The inset at the bottom shows the LC probabilistic template (blue) and the significant activation detected within this mask (red). The legend shows the t-values associated with color maps. This figure is adapted from ^11^.

Younger participants completed the fMRI session immediately following the baseline night but older participants were initially part of a different study ^24,25^ and completed the sMRI and fMRI recordings in addition to their initial engagement. Older participants completed the habituation and baseline night EEG recordings as part of their initial study and the sMRI and fMRI sessions were completed about 1.25y later as part of the current study (mean ±SD: 15.5 ±5.3 months). Prior to the fMRI session, older participants slept regularly for 1 week (verified with a sleep diary; based on our experience, actigraphy reports and sleep diaries do not deviate substantially in older individuals). Older participants were maintained in dim light (<10 lux) for 45min before the fMRI scanning.

### Sleep EEG metrics

Eleven channels were used for the baseline night (F3,z,4; C3,z,4; P3,z,4; O1,2) initially referenced to the left mastoid prior to re-referencing offline to the average of both mastoids (N7000 amplifier, EMBLA, Natus, Middleton, WI). Arousals and artefacts were detected automatically^26^ to provide the number of arousals during REM sleep, and excluded from the power spectral density analyses. Only frontal electrodes were considered in the analyses because the frontal region is most sensitive to sleep pressure manipulations ^27^; focusing on the frontal electrodes may also facilitate interpretation of future large-scale studies using ambulatory EEG, often restricted to frontal electrodes.

Sleep was staged in 30s-epochs using an automatic algorithm (ASEEGA, PHYSIP, Paris) ^28^ to provide total sleep time (TST). Averaged power was computed for each 30-minute bin, adjusted for the proportion of rejected data. The adjusted values were then summed across REM sleep^29^ to provide REM theta energy (overnight cumulated 4.25-8Hz power). Power in the other typical bands of the sleep EEG were computed similarly during both REM and NREM for specificity assessments (Delta band: 0.5-4Hz; Theta: 4.25-8Hz; Alpha: 8.25-12HZ; Sigma: 12.25-16Hz; Beta: 16.25-30Hz). Sigma power (12.25-16Hz) was computed during the 1-min preceding each REM episode, as the weighted sum of 4s artefact-free window (2s overlap per 30s epoch), prior to averaging over the number of REM episodes.

### Cognitive tasks

#### Visual perceptual rivalry task

The task (∼12min total duration) consisted of watching a 3D Necker cube, which can be perceived in two different orientations (**Figure 1B**), for 10 blocks of 1min separated by 10s of screen-center cross fixation. Participants were instructed to report switches between the two percepts through a button press.

#### Auditory salience detection task

The task (∼10min total duration) consisted of an oddball paradigm requiring reports on the perception of rare deviant target tones (1,000Hz, 100ms, 20% of tones) that were pseudo-randomly interleaved within a stream of standard stimuli (500Hz, 100ms) through a button press (**Figure 1C**). The task included 270 stimuli (54 targets).

These tasks were selected because they were thought to activate the LC. As we previously reported both tasks successfully triggered a response of the LC ^10,11^ **(Figure 1D and E)**.

### MRI data acquisition, preprocessing and univariate analyses

MRI data were acquired using a MAGNETOM Terra 7T MRI system (Siemens Healthineers, Erlangen, Germany), with a single-channel transmit and 32-receiving channel head coil (1TX/32RX, Nova Medical, Forchheim, Germany). Blood-oxygen-level-dependent (BOLD) fMRI data were acquired using a multi-band (MB) gradient-recalled echo–echo-planar imaging (GRE-EPI) sequence (main parameters: repetition time = 2.340ms, flip angle = 90°, 86 axial 1.4 mm–thick slices, no interslice gap, matrix size = 160 × 160, voxel size = 1.4 × 1.4 × 1.4 mm^3^,MB acceleration factor = 2, GeneRalized Autocalibrating Partial Parallel Acquisition (GRAPPA) acceleration factor = 3). The cardiac pulse and the respiratory movements were recorded concomitantly using, respectively, a pulse oximeter and a breathing belt (Siemens Healthineers). The fMRI acquisition was followed by a dual-echo 2D GRE field mapping sequence to assess B0 magnetic field inhomogeneities with the following parameters: TR = 5.2 ms, TEs = 2.26 ms and 3.28 ms, flip angle (FA) = 15°, bandwidth = 737 Hz/pixel, matrix size = 96 × 128, 96 axial slices with 2 mm thickness, voxel size = 2 × 2 × 2 mm ^3^, acquisition time = 1:38 minutes.

A Magnetization-Prepared with 2 RApid Gradient Echoes (MP2RAGE) sequence was used to acquire T1 anatomical images: TR = 4,300 ms, TE = 1.98 ms, FA = 5°/6°, TI = 940 ms/2,830 ms, bandwidth = 240 Hz/pixel, matrix size = 256 × 256, 224 axial 0.75 mm–thick slices, GeneRalized Autocalibrating Partial Parallel Acquisition (GRAPPA) acceleration factor = 3, voxel size = 0.75 × 0.75 × 0.75 mm^3^, acquisition time = 9:03 minutes.^30^

The LC-specific sequence consisted of a 3D high-resolution magnetization transfer– weighted turbo-flash (MT-TFL) sequence with the following parameters^31^: TR = 400 ms, TE = 2.55 ms, FA = 8°, bandwidth = 300 Hz/pixel, matrix size = 480 × 480 × 60, number of averages = 2, turbo factor = 54, magnetization transfer contrast (MTC) pulses = 20, MTC FA = 260°, MTC RF duration = 10,000 μs, MTC inter-RF delay = 4,000 μs, MTC offset = 2,000 Hz, voxel size = .4 × .4 × .5 mm^3^, acquisition time = 8:13 minutes. Axial slices were acquired and centered for the acquisitions perpendicularly to the rhomboid fossa (i.e., the floor of the fourth ventricle located on the dorsal surface of the pons) ^31^.

Functional and anatomical MRI data were preprocessed using SPM12, ANTs and SynthStrip brain extraction tool ^32^, as fully described previously ^10,11^. The preprocessed data were resampled to a 1mm^3^ resolution. Individual statistical analyses consisted of a general linear model (GLM) including one regressor of interest, consisting of a switch in perception (perceptual rivalry) or target tone (salience detection) modeled as an event (convolved with the canonical hemodynamic response function - HRF). Participant movement parameters, respiration, and heart rate were used as covariates of no interest (physiological data of 4 volunteers were not available and therefore not included in their individual design matrices). The T1 structural whole-brain image was used to extract individual total intracranial volume (TIV) using CAT12 toolbox ^33^.

Individual LC masks were manually delineated by 2 experts based on LC-specific images (as in ^10^) and activity of left LC was extracted in each subpart (as LC responses were more prominent in the left LC in both tasks ^10,11^). The different nuclei of the hypothalamus do not offer a lot of contrast in MRI images such that they cannot be segmented using current approaches. We therefore used an automated segmentation algorithm to parcellate the hypothalamus into 5 subparts which encompass several nuclei - anterior-inferior, anterior-superior, posterior, inferior tubular and superior tubular ^34^ (see **Figure 2A** for the nuclei deemed to be included in each subpart). We then extracted the activity of these subparts within the left hypothalamus (i.e., mean value over each subpart of hypothalamus) during both tasks for each participant using the REX Toolbox (https://web.mit.edu/swg/software.htm).

**Figure 2.**
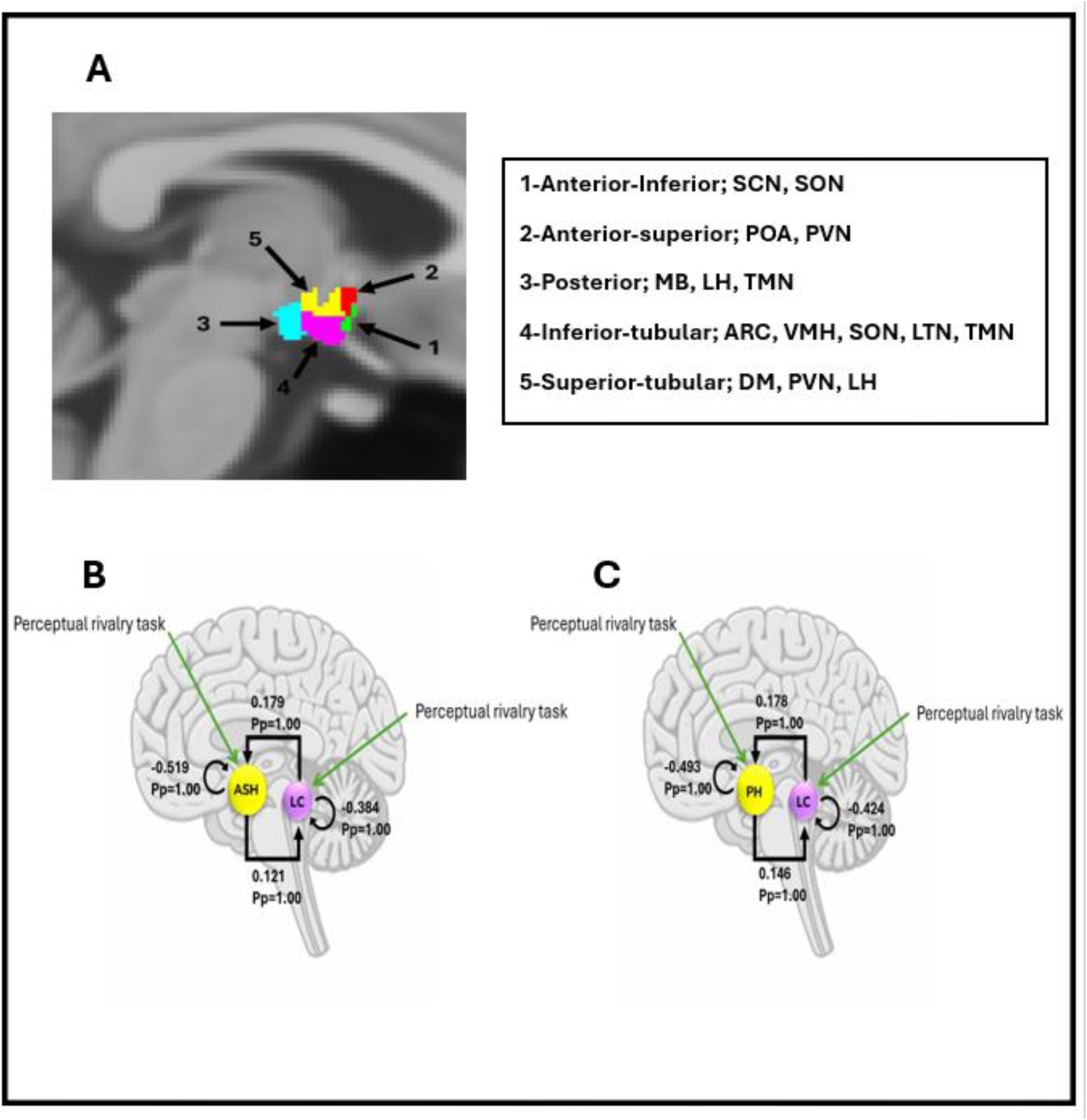
**>(A)** Segmentation of the hypothalamus in five subparts in a representative participant. The nuclei encompassed by the different subparts are indicated in the right inset – according to ^34^. ARC: arcuate nucleus; DMH; dorsomedial nucleus; LH lateral hypothalamus; LTN: lateral tubular nucleus; MB: mamillary body; POA: preoptic area; PVN: paraventricular nucleus; PNH: posterior nucleus of the hypothalamus; SCN: suprachiasmatic nucleus; SON: supraoptic nucleus; TMN: tuberomammillary nucleus; VMN: ventromedial nucleus. **(B)** The DCM model, which included intrinsic connections between anterior superior hypothalamus and locus coeruleus, along with self-feedback gain control connections for both regions. Task inputs were further considered to reach both regions. The DCM analysis showed that there was very strong evidence (Pp=1.0) for reciprocal positive influence between LC and anterior superior hypothalamus as well as self-inhibition in both LC and anterior superior hypothalamus. **(C)** The DCM model, which included intrinsic connections between posterior hypothalamus and locus coeruleus, along with self-feedback gain control connections for both regions. Task inputs were further considered to reach both regions. The DCM analysis showed that there was very strong evidence (Pp=1.0) for reciprocal positive influence between LC and posterior hypothalamus as well as self-inhibition in both LC and posterior hypothalamus.

### Effective Connectivity Analysis

Our previous analysis of the fMRI data of both tasks indicated that only a weak recruitment would be detected within the hypothalamus (i.e. no responses associated to the events of interest were detected using a relatively liberal p < .001 significance threshold). This does not, in principle, prohibit effective connectivity from being computed and importantly to be associated with other features of interest, such as sleep electrophysiology metrics. To select the hypothalamus subpart to be considered in our connectivity analyses, we reasoned that the activity of a given hypothalamus subpart should at least be weakly associated with our sleep metrics of interest. We therefore determined whether the activity estimate of each subpart – separately for each task - was associated with either REM theta energy or sigma power prior to REM episodes. Left hypothalamus subpart activity was used in Generalized Linear Mixed Models (GLMM) to seek correlation with REM theta energy and sigma power prior to REM sleep episodes (one GLMM per task and per sleep metric) (see statistic section below). Only the hypothalamus subparts that yielded at least a statistical trend (p < 0.1) with either sleep metrics during a given task were considered for effective connectivity analyses.

Dynamic Causal Modelling (DCM) framework^35^, implemented in SPM12, was used to compute the effective connectivity between the LC and each of the selected hypothalamus subparts during the selected task (i.e. the anterior-superior and posterior subparts during the Visual perceptual rivalry task – see results). BOLD signal time series were extracted from individually defined ROIs (i.e., LC and hypothalamus subparts masks), based on the individual statistical maps thresholded at p<.05 uncorrected, and used to infer the underlying neural activity. We extracted the first principal component (eigenvariate) of the "adjusted" time series, which represents the time series after regressing out effects of no interest, using the approach outlined by ^36^.

Stochastic DCM was used as we did not detect significant responses in the selected subparts of hypothalamus when inspecting the whole-brain statistical analyses over the entire sample (cf. above) ^37^. The DCM model included both intrinsic connections between the two regions, along with self-feedback gain control connections for LC and hypothalamus subpart (**Figure 2B and C**). Task inputs were further considered to reach both regions. Time series extracted from individual ROIs were subjected to a first-level DCM analysis, where the model was estimated for each subject. To isolate the connectivity parameters that were contributing to the model and could therefore be used in a GLMM seeking associations with sleep metrics (see below), we performed a Parametric Empirical Bayes (PEB) analysis ^38,39^ over the first-level DCM parameter estimates. PEB is a hierarchical Bayesian model that evaluates commonalities and differences among subjects in the effective connectivity domain at the group level by performing Bayesian model reduction (BMR), which explores the space of DCM models and leads to a subset of models best explaining the data and Bayesian model averaging (BMA) of the parameters across models weighted by the evidence of each model. Subsequently, as there is no concept of significance in Bayesian analysis, we reported and used in the GLMM, the parameters contributed to the model evidence with a posterior probability (Pp) exceeding 0.90.

### Statistics

GLMMs were performed in SAS 9.4 (SAS Institute, NC, USA) and were adjusted for the distribution of the dependent variables. Outliers among connectivity and sleep metrics lying beyond four times the standard deviation were removed from the analysis (maximum one data point was removed, the final number of individuals included in each analysis is reported in each table). The first GLMMs were meant to isolate which hypothalamus subparts would be included in DCM. They included the 2 sleep features of interest as dependent variables in each task separately and hypothalamus subpart, hypothalamus activity estimates and age-group, including sex, TST and TIV as covariates (i.e. 4 models in total, 1 per sleep metric and task). Our initial models included the three-way interaction between hypothalamus activity, hypothalamus subpart and age group and all three simple two-way interaction terms. Non-significant three-way / two-way interactions were removed from the models based on Bayesian Index Criterion (BIC) for fit quality estimation such that final models only included the hypothalamus activity and hypothalamus subpart interaction. This yielded statistical trends with the activity of the anterior superior and posterior subparts during the perceptual rivalry task (see results) for REM theta energy so that DCM included these 2 subparts during the latter task.

The next set of GLMMs tested for associations between connectivity metrics and REM theta energy, again used as dependent variable, and including sex, TST and TIV as covariates. Following the same procedure described in the preceding paragraph, all final models included an interaction term between the connectivity metric and the age-group. Semi-partial R^2^ (R^2*^) values were computed to estimate the effect sizes of significant fixed effects and statistical trends in all GLMMs ^40^. Significance was determined following the Benjamini-Hochberg procedure for False discovery rate procedure [p<.025 (for rank 1/2); p<.05 (for rank 2/2)].

We computed a prior sensitivity analysis to get an indication of the minimum detectable effect size in our main analyses given our sample size. According to G*Power 3 (version 3.1.9.4) ^41^; taking into account a power of .8, an error rate α=.025, and a sample size of 51, we could detect medium effect sizes *r*>.39 (2-sided; CI:.13–.6; *R*²>.15, CI:.02–.36) within a linear multiple-regression framework including 2 tested predictor (connectivity, age group) and 2/3 covariates (sex, TIV, TST where relevant).

## Results

Fifty-one healthy individuals aged 18 to 31y (N=33; 27 women) and 50 to 70y (N=18; 14 women) completed an fMRI protocol. We first selected the hypothalamus subpart to include in our connectivity analyses in each task (see method). Statistical analyses consisted of two GLMMs per task respectively including REM theta energy and sigma power prior to REM sleep as the dependent variables. Considering the perceptual rivalry task, we found a statistical trend for the association between REM theta energy and the interaction between the activity of the five subparts of the hypothalamus and hypothalamus subpart (f=2.05; p=.08; Table 2). Post hoc contrasts revealed that higher REM theta energy shows a statistical trend with a lower response of the anterior-superior hypothalamus (t= -1.81; p=.07) and higher response of the posterior hypothalamus (t=1.90; p=.05; **Figure 3A and B**) while other subparts activity did not show such a trend (t< -0.02 ;p> 0.18) (**Suppl. Figure S1, Suppl. Table S1**). The GLMM with the sigma power prior to REM and hypothalamus subpart activity in the perceptual rivalry task did not reveal any associations. **(Suppl. Figure S2, Table 2)**. Considering the auditory salience detection task, the GLMM with either sleep metrics did not reveal any association with hypothalamus subpart activity, even as a statistical trend. **(Suppl. Figure S3 and S4, Table 2)**.

**Figure 3.**
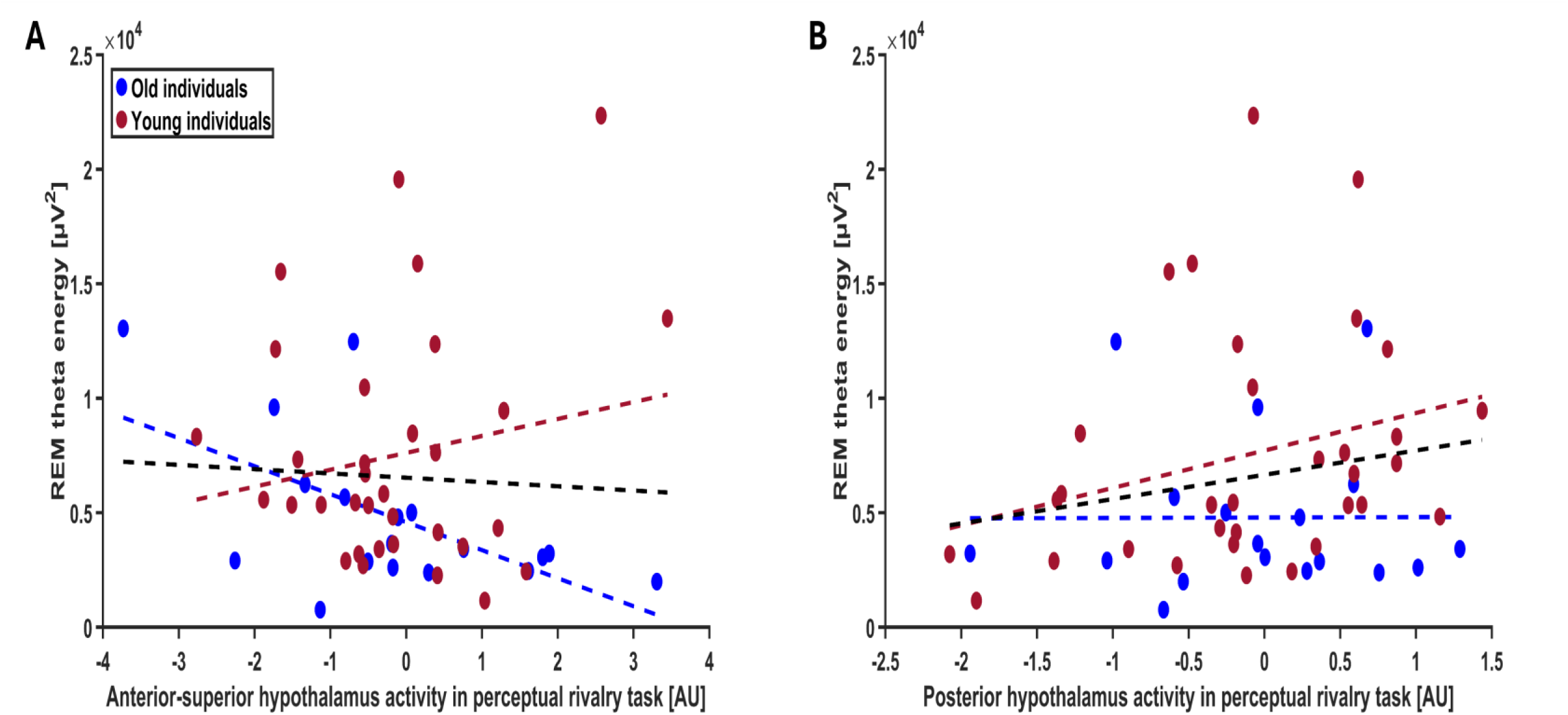
**(A)** Association between REM theta energy and the anterior-superior hypothalamus activity during the perceptual rivalry task. The GLMM yielded to a statistical trend for the hypothalamus activity by hypothalamus subpart interaction (p=0.08), and post hoc analyses led to a statistical trend for the negative association of the anterior-superior hypothalamus activity (p=0.07). **(B)** Association between REM theta energy and the posterior hypothalamus activity during the perceptual rivalry task. The GLMM yielded to a statistical trend for the hypothalamus activity by hypothalamus subpart interaction (p=0.08), and post hoc analyses led to a statistical trend for the negative association of the posterior hypothalamus activity (p=0.05). Simple regression lines are used for a visual display and do not substitute the GLMM outputs (Table 2). The black line represents the regression irrespective of age groups (young + old). Non-significant associations between REM theta energy as well as sigma power prior to REMS and the activity of subparts of hypothalamus during the perceptual rivalry task and salience detection task are displayed on Suppl. Figure S1-S4.

**Table 2.** The association between 2 sleep metrics of interest and 5 hypothalamus subparts activity via the perceptual rivalry task and silence detection task.

| Task | Sleep metric (dependent variable) | hypothalamus activity | hypothalamus subpart | hypothalamus activity * hypothalamus subpart | Age group | Sex | TIV | Total sleep time |
| --- | --- | --- | --- | --- | --- | --- | --- | --- |
| perceptual rivalry task | REM Theta energy (N=51) | F(1,239)=0.62<br>P=0.432 | F(4,239)=0.05<br>P=0.994 | F(4,239)=2.05<br>**P=0.088**<br>**R<sup>2</sup>=0.033** | F(1,239)=0.44<br>P=0.507 | F(1,239)=0.94<br>P=0.334 | F(1,239)=0.53<br>P=0.466 | F(1,239)=28.5<br>**P<0.0001**<br>**R<sup>2</sup>=0.106** |
| perceptual rivalry task | Sigma power prior to REMS (N=51) | F(1,239)=0.58<br>P=0.446 | F(4,239)=0.0<br>P=1.00 | F(4,239)=0.32<br>P=0.864 | F(1,239)=2.74<br>P=0.098 | F(1,239)=20.19<br>**P<0.0001**<br>**R<sup>2</sup>=0.077** | F(1,239)=15.03<br>**P=0.0001**<br>**R<sup>2</sup>=0.059** | F(1,239)=1.52<br>P=0.218 |
| silence detection task | REM Theta energy (N=51) | F(1,239)=0.0<br>P=0.950 | F(4,239)=0.01<br>P=0.999 | F(4,239)=0.22<br>P=0.927 | F(1,239)=1.28<br>P=0.258 | F(1,239)=1.70<br>P=0.193 | F(1,239)=3.79<br>P=0.052 | F(1,239)=33.00<br>**P<0.0001**<br>**R<sup>2</sup>=0.121** |
| silence detection task | Sigma power prior to REMS (N=51) | F(1,234)=5.0<br>**P=0.026**<br>**R<sup>2</sup>=0.020** | F(4,234)=0.01<br>P=0.999 | F(4,234)=0.65<br>P=0.628 | F(1,234)=3.74<br>P=0.054 | F(1,234)=24.94<br>**P<0.0001**<br>**R<sup>2</sup>=0.096** | F(1,234)=14.62<br>**P=0.0002**<br>**R<sup>2</sup>=0.058** | F(1,234)=2.28<br>P=0.132 |

Prior to the analysis, we removed the outliers among connectivity and sleep metrics by excluding the samples lying beyond four times the standard deviation (the final number of individuals included in each analysis is reported below each dependent variable).
REM: rapid eye movement; REMS: rapid eye movement sleep.

Therefore, the DCM analysis concentrated on the anterior-superior and posterior subparts of the hypothalamus during the perceptual rivalry task and sought relationship with REM theta energy, as it was the sole sleep metric that demonstrated a statistical trend with hypothalamus activity. The DCM analysis first showed that there was very strong evidence for reciprocal positive influence between LC and both subparts of hypothalamus (Pp=1.0; **Figure 2B and C**), confirming that the connectivity anticipated based on animal model ^2^ can be detected in humans using a stochastic DCM approach.

Considering the connectivity from anterior-superior subpart of hypothalamus to LC, the GLMM revealed a significant connectivity-by-age-group interaction when examining REM theta energy (t=2.30; p=0.026), on top of a main effect of TST, while the other covariates were not significant **(Table 3)**. Post hoc contrasts highlighted that higher connectivity was associated with lower REM theta energy in the older (t=-2.09; p=0.042) but not in the younger group (t=1.06; p=0.294) **(Figure 4A)**. In addition, if a participant putatively outlier for TIV is removed (SD=4.078), the connectivity-by-age-group interaction became even more robust (t=2.37; p=0.022; i.e. below the multiple comparison correction p < 0.025 threshold) with post hoc contrasts identically showing the association between higher connectivity and lower REM theta energy in the older group (t = -2.12, p = 0.039), but not in the younger group (t = 1.15, p = 0.255). We then considered the connectivity from LC to anterior-superior subpart of hypothalamus, but GLMM did not yield significant association with REM theta energy **(Table 3**; **Figure 4B)**. Likewise, when we considered the connectivity between the posterior hypothalamus subpart and the LC (i.e. both to and from the LC), GLMMs did not lead to any significant association with REM theta energy **(Table 3**; **Figure 4C and D)**.

**Figure 4.**
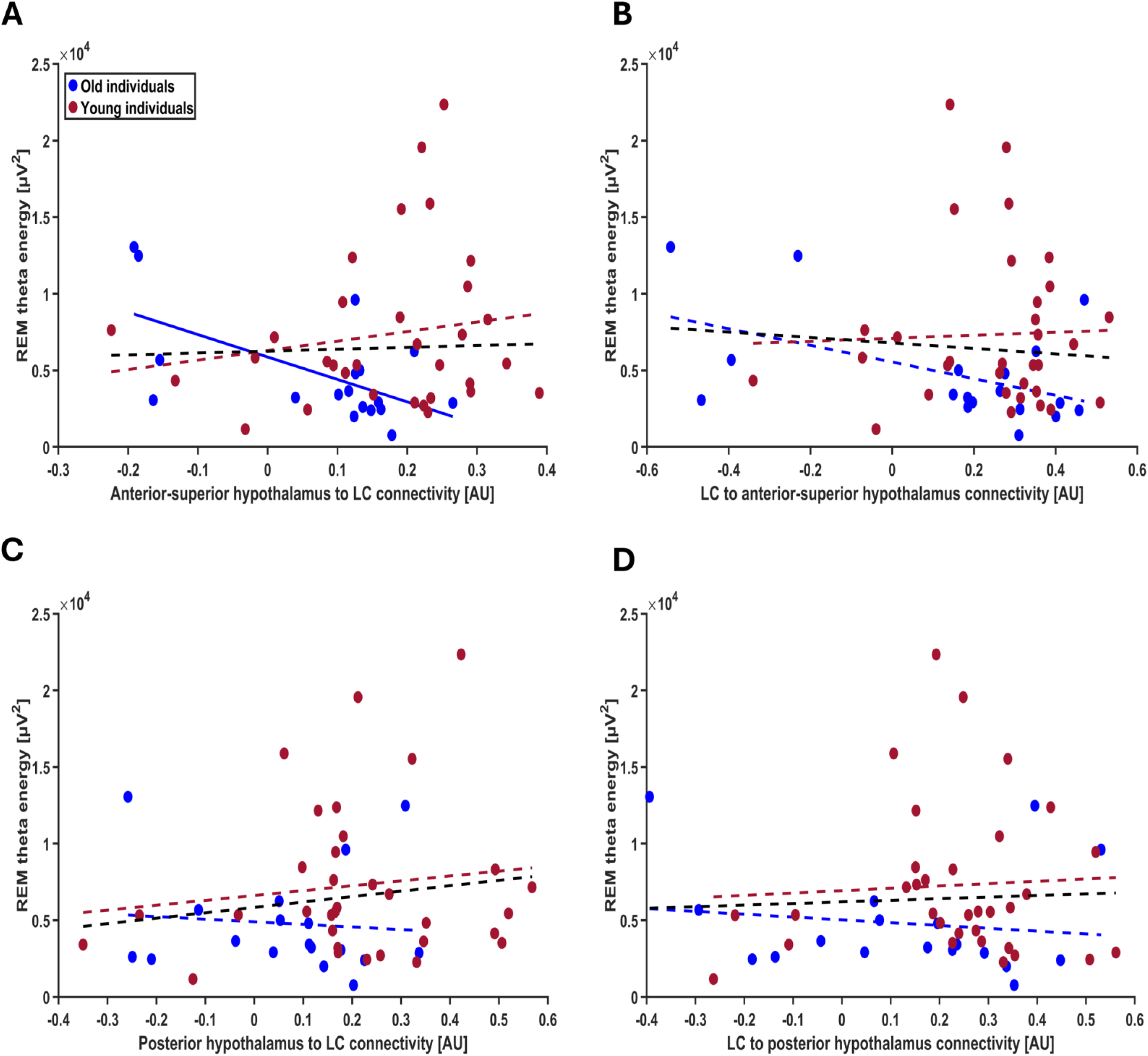
**(A)** Association between REM theta energy and the anterior-superior hypothalamus to LC connectivity. The GLMM yielded to a significant age group by connectivity interaction (p=0.026), and post hoc analyses led to a significant association for the older (p=0.042) but not the young group (p=0.294). After excluding one putative outlier in TIV (≥4 SD), the connectivity-by-age-group interaction became even more robust (t=2.37; p=0.022) with post hoc contrasts reaffirming the association between higher connectivity and lower REM theta energy in the older group (t = -2.12, p = 0.039), but not in the younger group (t = 1.15, p = 0.255). **(B)** Association between REM theta energy and LC to anterior-superior hypothalamus connectivity. The GLMM did not show a significant age group by connectivity interaction (p=0.144). **(C)** Associations between REM theta energy and the connectivity from posterior hypothalamus to LC. The GLMM did not show a significant age group by connectivity interaction (p=0.326). **(D)** Association between REM theta energy and the connectivity from LC to posterior hypothalamus. The GLMM did not show a significant age group by connectivity interaction (p=0.439). Simple regression lines are used for a visual display and do not substitute the GLMM outputs (Table 3). The black line represents the regression irrespective of age groups. Solid and dashed regression lines represent significant and non-significant outputs of the GLMM, respectively.

**Table 3.** Associations between REM Theta energy and the connectivity between anterior-superior hypothalamus and LC as well as between posterior hypothalamus and LC.

| Type of connectivity | Sleep metric (dependent variable) | connectivity | Age group | connectivity*age group | Sex | TIV | Total sleep time |
| --- | --- | --- | --- | --- | --- | --- | --- |
| From anterior-superior hypothalamus to LC | REM Theta energy (N=50) | F(1,43)=1.09<br>P=0.302 | F(1,43)=0.62<br>P=0.436 | F(1,43)=5.29<br>**P=0.026<sup>A</sup>**<br>**R<sup>2</sup>=0.109** | F(1,43)=0.02<br>P=0.902 | F(1,43)=0.03<br>P=0.854 | F(1,43)=4.74<br>**P=0.035**<br>**R<sup>2</sup>=0.099** |
| From LC to anterior-superior hypothalamus | REM Theta energy (N=50) | F(1,43)=0.38<br>P=0.539 | F(1,43)=0.17<br>P=0.686 | F(1,43)=2.21<br>P=0.144 | F(1,43)=0.05<br>P=0.825 | F(1,43)=0.00<br>P=0.981 | F(1,43)=4.73<br>**P=0.035**<br>**R<sup>2</sup>=0.099** |
| From posterior hypothalamus to LC | REM Theta energy (N=50) | F(1,43)=0.00<br>P=0.994 | F(1,43)=0.09<br>P=0.765 | F(1,43)=0.98<br>P=0.326 | F(1,43)=0.58<br>P=0.451 | F(1,43)=0.12<br>P=0.726 | F(1,43)=4.77<br>**P=0.034**<br>**R<sup>2</sup>=0.099** |
| From LC to posterior hypothalamus | REM Theta energy (N=50) | F(1,43)=0.14<br>P=0.710 | F(1,43)=0.01<br>P=0.911 | F(1,43)=0.61<br>P=0.439 | F(1,43)=0.49<br>P=0.488 | F(1,43)=0.08<br>P=0.776 | F(1,43)=4.40<br>**P=0.041**<br>**R<sup>2</sup>=0.092** |

Prior to the analysis, we removed the outliers among connectivity and sleep metrics by excluding the samples lying beyond four times the standard deviation (the final number of individuals included in each analysis is reported below each dependent variable).
<sup>A</sup> p = 0.022 and F(1,42)=5.61 after excluding one putative outlier in TIV ( $\geq 4$ SD).
LC: locus coeruleus; TIV: total intracranial volume; REM: rapid eye movement; REMS: rapid eye movement sleep.

In the next steps, we verified the specificity of our finding for REM theta energy and turned toward the other frequency bands of the EEG during both REM and NREM and found that although association did not extend to all frequency bands, they were not restricted to REM theta energy. Separate GLMMs found that connectivity from the anterior-superior subpart of the hypothalamus to the LC by age group interaction was significantly associated with several lower frequency bands of both REM and NREM: interaction was significant for alpha energy in REMS (t=3.21; p=0.0025), delta energy in NREMS (t=2.51; p=0.016) theta energy in NREMS (t=2.91; p=0.006) and alpha energy in NREM (t=3.17; p=0.0028) but not for delta, sigma and beta energy in REM (t< 2.03; p >0.048) and sigma and beta energy in NREM (t < 0.60; p > 0.550); each time we observed a significant negative association between connectivity and frequency band energy in the older group (REM alpha: t=-2.79; p=0.008; NREM delta: t=-2.04; p=0.047; NREM theta: t=-2.34; p=0.024; alpha NREM: t=-2.75; p=0.009) but not younger group (REM alpha: t=1.64; p=0.11; NREM delta: t=1.49; p=0.14; NREM theta: t=1.76; p=0.085; alpha NREM: t=1.68; p=0.099) **(Figure 5A, B, C and D, Table 4)**.

**Figure 5.**
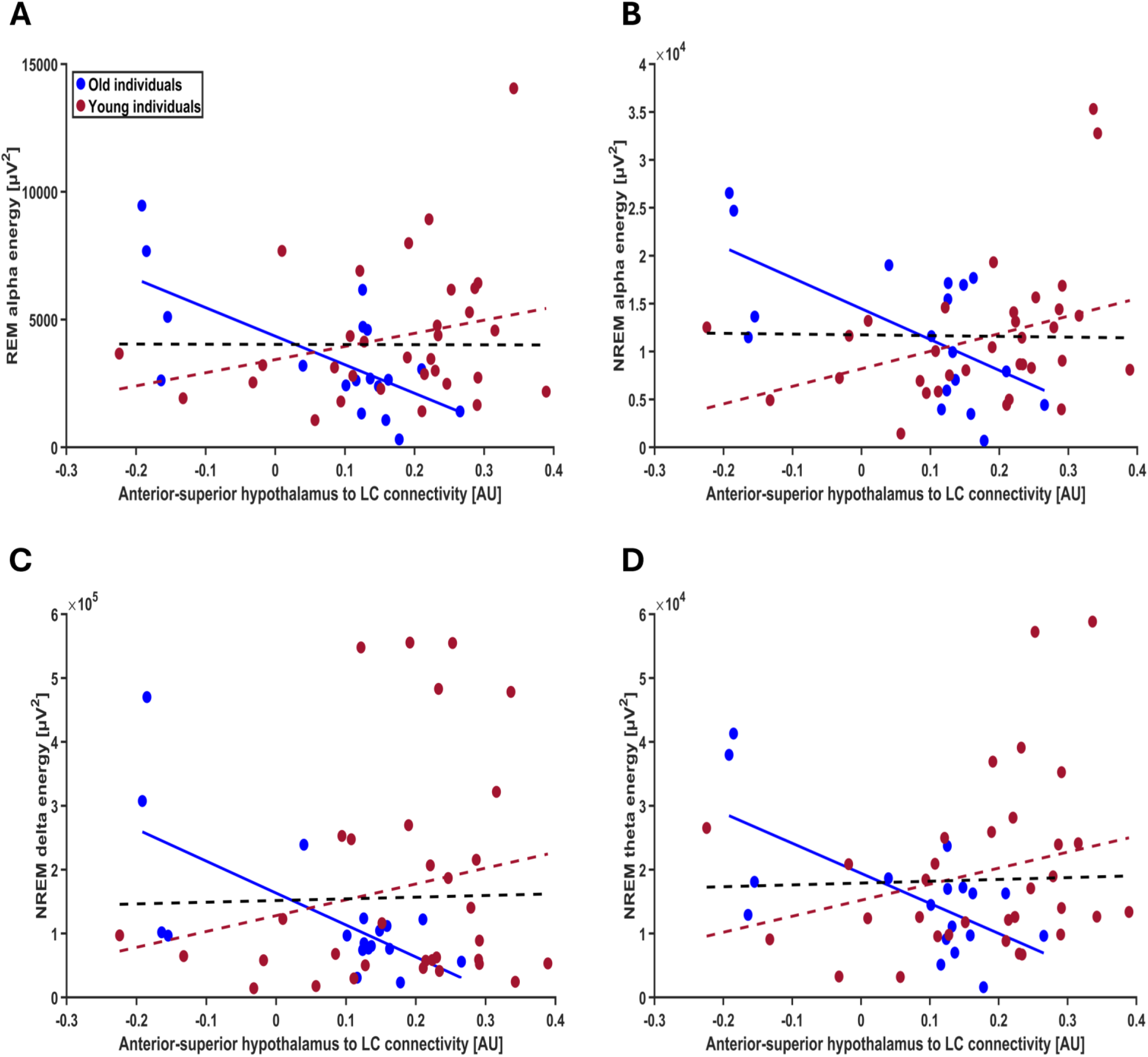
**(A)** Association between REM alpha energy and the anterior-superior hypothalamus to LC connectivity. The GLMM yielded a significant age group by connectivity interaction (p=0.002), and post hoc analyses led to a significant association for the older (p=0.007) but not the younger group (p=0.108). **(B)** Association between NREM alpha energy and the anterior-superior hypothalamus to LC connectivity. The GLMM yielded a significant age group by connectivity interaction (p=0.002), and post hoc analyses led to a significant association for the older (p=0.0008) but not the younger group (p=0.099). **(C)** Association between NREM delta energy and the anterior-superior hypothalamus to LC connectivity. The GLMM yielded a significant age group by connectivity interaction (p=0.015), and post hoc analyses led to a significant association for the older (p=0.047) but not the younger group (p=0.142). **(D)** Association between NREM theta energy and the anterior-superior hypothalamus to LC connectivity. The GLMM yielded a significant age group by connectivity interaction (p=0.005), and post hoc analyses led to a significant association for the older (p=0.023) but not the younger group (p=0.085). Simple regression lines are used for a visual display and do not substitute the GLMM outputs (Table 4). The black line represents the regression irrespective of age groups. Solid and dashed regression lines represent significant and non-significant outputs of the GLMM, respectively. Non-significant associations between the energy of other frequency bands and the connectivity from anterior-superior hypothalamus to LC to test the specificity are displayed on Suppl. Figure S5.

**Table 4.** Associations between exploratory sleep metrics and the connectivity from anterior-superior hypothalamus to LC.

| Sleep metric (dependent variable) | connectivity | Age group | connectivity*age group | Sex | TIV | Total sleep time |
| --- | --- | --- | --- | --- | --- | --- |
| REM delta energy (N=51) | F(1,44)=0.49<br>P=0.487 | F(1,44)=0.66<br>P=0.420 | F(1,44)=4.11<br>**P=0.048\***<br>**R<sup>2</sup>=0.085** | F(1,44)=0.11<br>P=0.745 | F(1,44)=0.13<br>P=0.718 | F(1,44)=0.86<br>P=0.358 |
| REM alpha energy (N=50) | F(1,43)=1.74<br>P=0.194 | F(1,43)=2.92<br>P=0.094 | F(1,43)=10.29<br>**P=0.002**<br>**R<sup>2</sup>=0.193** | F(1,43)=0.04<br>P=0.851 | F(1,43)=0.10<br>P=0.757 | F(1,43)=5.43<br>**P=0.024**<br>**R<sup>2</sup>=0.112** |
| REM sigma energy (N=51) | F(1,44)=0.00<br>P=0.963 | F(1,44)=0.83<br>P=0.366 | F(1,44)=0.07<br>P=0.795 | F(1,44)=0.46<br>P=0.503 | F(1,44)=0.26<br>P=0.611 | F(1,44)=6.08<br>**P=0.017**<br>**R<sup>2</sup>=0.121** |
| REM beta energy (N=50) | F(1,43)=0.15<br>P=0.703 | F(1,43)=0.11<br>P=0.743 | F(1,43)=0.82<br>P=0.369 | F(1,43)=5.50<br>**P=0.023**<br>**R<sup>2</sup>=0.113** | F(1,43)=4.19<br>**P=0.046**<br>**R<sup>2</sup>=0.088** | F(1,43)=12.02<br>**P=0.001**<br>**R<sup>2</sup>=0.221** |
| NREM delta energy (N=51) | F(1,44)=0.59<br>P=0.445 | F(1,44)=2.02<br>P=0.162 | **F(1,44)=6.31**<br>**P=0.015**<br>**R<sup>2</sup>=0.125** | F(1,44)=0.13<br>P=0.722 | F(1,44)=1.30<br>P=0.260 | F(1,44)=0.22<br>P=0.642 |
| NREM theta energy (N=51) | F(1,44)=0.73<br>P=0.396 | F(1,44)=2.21<br>P=0.144 | F(1,44)=8.48<br>**P=0.005**<br>**R<sup>2</sup>=0.161** | F(1,44)=0.14<br>P=0.714 | F(1,44)=0.57<br>P=0.452 | F(1,44)=1.33<br>P=0.254 |
| NREM alpha energy (N=51) | F(1,44)=0.83<br>P=0.368 | F(1,44)=4.36<br>**P=0.042**<br>**R<sup>2</sup>=0.090** | F(1,44)=10.04<br>**P=0.002**<br>**R<sup>2</sup>=0.185** | F(1,44)=0.09<br>P=0.765 | F(1,44)=0.00<br>P=0.973 | F(1,44)=0.37<br>P=0.546 |
| NREM sigma energy (N=51) | F(1,44)=0.35<br>P=0.559 | F(1,44)=0.00<br>P=0.953 | F(1,44)=0.32<br>P=0.575 | F(1,44)=0.02<br>P=0.899 | F(1,44)=0.13<br>P=0.717 | F(1,44)=3.79<br>P=0.058 |
| NREM beta energy (N=51) | F(1,44)=0.17<br>P=0.680 | F(1,44)=0.75<br>P=0.390 | F(1,44)=0.36<br>P=0.550 | F(1,44)=0.06<br>P=0.804 | F(1,44)=0.20<br>P=0.657 | F(1,44)=4.16<br>**P=0.047**<br>**R<sup>2</sup>=0.086** |

Prior to the analysis, we removed the outliers among connectivity and sleep metrics by excluding the samples lying beyond four times the standard deviation (the final number of individuals included in each analysis is reported below each dependent variable).
\*post hoc analysis showed that the significant association between REM delta energy and age group by connectivity interaction considering the connectivity from anterior-superior hypothalamus to LC (p=0.048) is driven by the association between REM delta energy and the difference of connectivity values between two age groups and not each group separately.
LC: locus coeruleus; TIV: total intracranial volume; REM: rapid eye movement; REMS: rapid eye movement sleep; NREM: non-rapid eye movement.

## Discussion

The interplays between the subcortical structures regulating sleep are not established in humans. Here, we used 7 Tesla fMRI to capture the crosstalk between the hypothalamus and the locus coeruleus and related it to the electrophysiology of REM sleep. We presumed that the connectivity between subparts of the hypothalamus and the LC during wakefulness would reflect in part their connectivity during sleep. We provide evidence that lower REM theta energy is associated with higher effective connectivity from the anterior-superior hypothalamus, which encompasses the preoptic area, to LC in the older individuals of our sample. The association was not specific to REM theta energy and extended to other (but not all) lower frequency bands of both REM and NREM sleep. These findings constitute an original investigation of how a small network of subcortical areas may take part in sleep regulation in humans and provide novel insights into the changes in sleep taking place over the healthy lifespan.

The tasks included in the protocol were geared toward ensuring a reliable recruitment of the LC ^10,11^ while they did not strongly recruit the hypothalamus. We therefore used an effective connectivity approach that was applicable to such cases (i.e. stochastic DCM). We further isolated candidate hypothalamus subparts that could be included in our connectivity models by seeking at least weak associations with our two sleep metrics of interest. We found weak indications that the activity of the posterior and anterior-superior subparts of the hypothalamus during the perceptual rivalry task were associated with REM theta energy, but not with sigma power prior to REM episodes. These indications were used to guide our connectivity analyses and will not be interpreted further although they warrant future investigations. We stress that the fact that we focused on specific subparts of the hypothalamus does not preclude the connectivity between the LC and other subparts of the hypothalamus that would be assessed in other contexts to be related to sleep electrophysiology, e.g. the anterior-inferior subpart encompassing the SCN ^15^ (i.e. if activity was assessed using different cognitive tasks, resting state fMRI, or a different vigilance state, etc.). Likewise, the connectivity of the LC with other parts of the brain in the perceptual rivalry as well as in the salience detection task may turn out to be related to sleep physiology in other analyses. The fact that we obtained very high evidence (Pp = 1) that the posterior and anterior-superior subparts were part of a network with the LC demonstrate that one can grasp a meaningful part of the complex interplays between the LC and nuclei of the hypothalamus *in vivo* in humans following our approach. There was indeed no guarantee that the network we constructed would be related to the fMRI signal we extracted for the hypothalamus and the LC. Our findings suggest that the mutual influence of the anterior-superior and posterior subparts of hypothalamus on the LC repeatedly demonstrated in animal ^42^ can be isolated in humans. This important proof-of-concept paves the way for future investigations that could be built around other regions and more complex networks.

Following the extraction of the connectivity metrics in the 2 networks respectively composed of the posterior subpart of the hypothalamus and the LC and of anterior-superior subparts of the hypothalamus and the LC, we find only one metric of the latter networks to be associated with REM theta energy. The anterior-superior subpart included the preoptic area, key to sleep regulation, but also the paraventricular nucleus (PVN) ^34^, which is typically related to food intake, energy balance ^43^, and vegetative regulation ^44^. Hence, we privilege that the associations that we found were mostly driven by preoptic area, which contains the ventrolateral (VLPO), lateral (LPO) and median (MPO) preoptic areas, which have all been involved in sleep regulation ^2,45,46^. The neurons of the preoptic area promote sleep onset and sleep maintenance by inhibitory GABAergic modulation of multiple arousal systems such as LC ^42^. The inhibitory action of the VLPO exerted on LC is considered as a requirement for sleep onset ^3^.

We find that the stronger the connectivity from the anterior-superior hypothalamus to the LC during wakefulness, the lower REM theta energy in older individuals. Theta oscillations consist of the most typical oscillatory mode of REM sleep. They are considered to be cortical correlates of the hippocampus ripple waves occurring during sleep and related to the memory function of REM sleep ^47^. We interpret these as a reflection of REM sleep intensity. We previously reported that a larger expression of LC responses during the same task during wakefulness was associated with a better expression of REM sleep theta oscillations ^11^. Our current finding may therefore indicate that a stronger connection between the preoptic nuclei and the LC prevents the LC from favoring REM sleep. The connectivity between the preoptic area and the LC, at least during wakefulness and potentially also during sleep, would contribute to REM sleep variability among older individuals and also, potentially, to the decreased expression of REM associated with aging. These potential actions may be related to the overall alteration in the balance of the neural circuits previously reported in aging ^48^. Alternatively, the impact of the preoptic nuclei onto the LC may prevent LC over reactivity that could become detrimental for the expression of REM sleep ^11^.

Interestingly, we found that the associations between anterior-superior hypothalamus to LC connectivity and neural oscillations extend beyond the theta energy in REMS. In older individuals, its negative correlations with alpha energy in REMS and NREMS as well as delta and theta energy in NREMS suggest a potential broader role for this connectivity in influencing sleep electrophysiology. Delta energy in NREMS (i.e., slow wave energy) is recognized as a marker of sleep need as well as sleep maintenance, quality, and restorative processes ^49^. Higher theta energy in NREMS is also linked to memory reactivation and memory consolidation ^50^. In addition, alpha power during REMS and NREMS is related to better cognitive performance as it is shown that individuals with cognitive impairment have lower alpha power in REMS and NREMS compared to healthy individuals ^51^. Similarly to our main link of interest with REM theta energy, the negative association between the strength of the connectivity between the preoptic nuclei of the hypothalamus and the LC may prevent the LC from favoring these brain oscillations during sleep. It could also represent an adaptive mechanism in older individuals to prevent over reaction of the LC. In any case it shows that the connectivity from the anterior-superior subpart of the hypothalamus to the LC is associated with neuronal synchrony over the lower range of the EEG spectral bands during sleep, potentially affecting the mechanism related to this synchrony (e.g. memory consolidation, sleep homeostasis). This could contribute to the previous findings on the aging brain’s adaptive modifications in homeostatic sleep control ^52^.

As noted earlier,^10,11^ while this research sheds light on the associations between hypothalamus-LC effective connectivity and variations in human sleep, certain limitations are inherent to the study. Young participants underwent fMRI scanning the day after their baseline night of sleep, while for the older group, there was an approximately one-year interval between the baseline sleep night and the fMRI session. Although sleep changes during the lifetime ^53^, it tends to remain stable over shorter periods (e.g. a few years) ^54^. Therefore, we believe this significant limitation does not fully account for our findings. Additionally, while we believe that wakefulness partly reflects brain activity and connectivity during sleep, future studies should examine these findings in the context of sleep to confirm it. Despite the extensive data collection involved, the sample size, particularly for the older cohort, is relatively small. To robustly validate our results, future studies should include larger, sex-balanced samples and uniform protocols across different age groups. Finally, our sample was predominantly female, a factor accounted for in our statistical analysis but still limiting the broad generalizability of the results.

In summary, we show that the connectivity between key subcortical structures for sleep regulation assessed during wakefulness may contribute to the variability of sleep electrophysiology. Our main finding includes the dominant oscillatory mode of REM, as a potential reflection of REM intensity and amnesic function. The association is detected in participants aged between 50 and 70y and beyond REM theta rhythms, suggesting a more prominent impact in the fragile sleep found in aging and on neuronal synchrony over lower frequencies. These results underscore the age-dependent modulation of LC circuitry and its implications for sleep regulation and stability. Understanding these neural dynamics provides new insights into the mechanisms underlying age-related sleep changes and may contribute to individually tailored future interventions aimed at improving sleep health across the lifespan.

## Supporting information

SUPPLEMENTARY MAERIAL

## Acknowledgments

This work was conducted at the GIGA-In Vivo Imaging platform of ULiège, Belgium. We thank Nikita Beliy, Alexandre Berger, Christian Berthomier, Islay Campbell, Ismael Dardour Hamzaoui, Erik Lambot, Catherine Hagelstein, Sophie Laloux, Annick Claes, Christian Degueldre, Brigitte Herbillon, Gregory Hammad, Patrick Hawotte, Thibault Gendron, Heidi I.L. Jacobs, Benjamin Lauricella, Alexia Lesoinne, Pierre Maquet, Thomas Pontus, John Read and Aubry Robert for their help in the different stages of the study. This work was supported by Fonds National de la Recherche Scientifique (FRS-FNRS, T.0242.19, T.0238.23 and J. 0222.20), Action de Recherche Concertée – Fédération Wallonie-Bruxelles (ARC SLEEPDEM 17/27-09), Fondation Recherche Alzheimer (SAO-FRA 2019/0025 & 2022/0014), Fondation Léon Fredericq, ULiège, European Regional Development Fund (Radiomed, Biomed-Hub, WALBIOIMAGING).

EB was supported by the Maastricht University - ULiège Imaging Valley. RS and FB were supported by the European Union’s Horizon 2020 research and innovation program under the Marie Skłodowska-Curie grant agreement no. 860613. RS is supported by ULiège. NM, IP, CP, EK, MZ, CB, FC, FB and GV are/were supported by the FRS-FNRS. PT and LL are/were supported by the EU Joint Programme Neurodegenerative Disease Research (JPND) IRONSLEEP and SCAIFIELD projects, respectively – FNRS references: PINT-MULTI R.8011.21 & 8006.20. LL is supported by the European Regional Development Fund (WALBIOIMAGING).

## Author contributions

Study concept and design by NM and GV. Data acquisition and analysis by NM, PT, EK, RS, EB, IP & FB. Methodological support and/or support in the interpreting the data by CP, FC, MZ, LL. Funding was mostly obtained by FC, CB, CP, and GV. NM and GV drafted the first version of the manuscript. All authors revised the manuscript and had final responsibility for the decision to submit for publication.

## Disclosure Statement

Financial Disclosure: none. Non-financial Disclosure: none.

## Data availability

The processed data and analysis scripts supporting the results included in this manuscript are publicly available via the following open repository: https://gitlab.uliege.be/CyclotronResearch-Centre/Public/fasst/xxx (to be defined upon acceptance of the paper). The raw data could be identified and linked to a single subject and represent a large amount of data and cannot be openly shared. Researchers willing to access to the raw data should send a request to the corresponding author (GV). Data sharing will require evaluation of the request by the local Research Ethics Board and the signature of a data transfer agreement (DTA).

## Notes

### Competing Interest Statement

The authors have declared no competing interest.

