## SUPPLEMENTARY MAERIAL for "The crosstalk between the anterior hypothalamus and the locus coeruleus during wakefulness is associated with low frequency oscillations power during sleep"

**Supplementary Table S1. Post hoc contrast on the associations between REMS theta energy and hypothalamus subparts activity estimated via the visual perceptual rivalry task.**

| Hypothalamus subpart | Estimate | DF | t value | P |
|---|---|---|---|---|
| **Interior-inferior hypothalamus** | -0.001 | 239 | -0.02 | 0.983 |
| **Anterior-superior hypothalamus** | -0.121 | 239 | -1.81 | **0.071** |
| **Posterior hypothalamus** | 0.194 | 239 | 1.90 | **0.058** |
| **Inferior-tubular hypothalamus** | -0.116 | 239 | -1.15 | 0.252 |
| **Superior-tubular hypothalamus** | -0.104 | 239 | -1.32 | 0.187 |

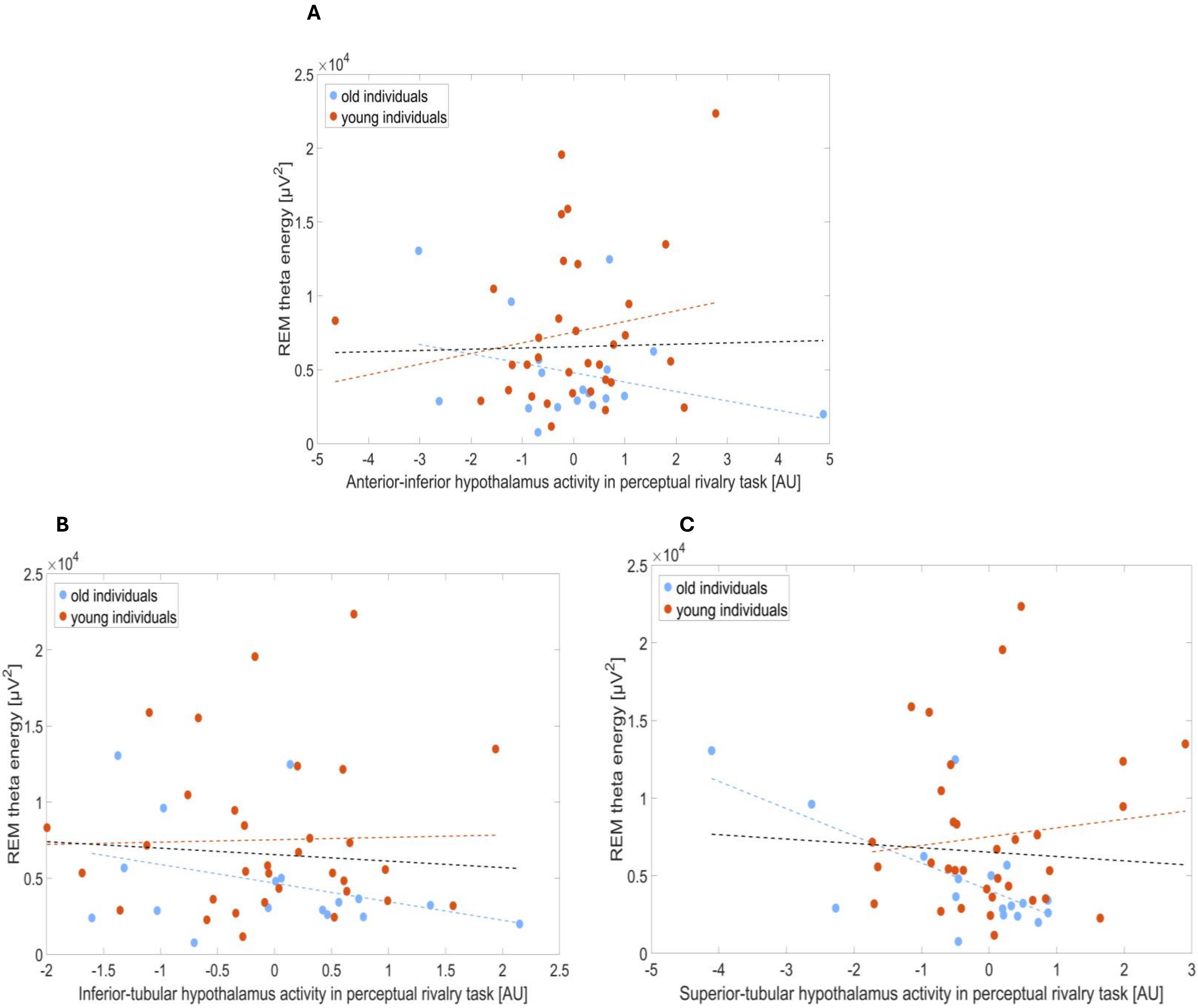


**Supplementary Figure S1. Non-significant associations between hypothalamus subparts activity estimates during the perceptual rivalry task and REM theta energy. (A)** Association between the interior-inferior hypothalamus activity estimates during the perceptual rivalry task and REM theta energy. **(B)** Association between the inferior-tubular hypothalamus activity estimates during the perceptual rivalry task and REM theta energy. **(C)** Association between the superior-tubular hypothalamus activity estimates during the perceptual rivalry task and REM theta energy.

Although the GLMM yielded to a statistical trend for the hypothalamus activity by hypothalamus subpart interaction (p=0.8), post hoc analyses did not show a statistical trend for any of these hypothalamus subparts (p>0.18).

Simple regression lines are used for a visual display and do not substitute the GLMM outputs. The black line represents the regression irrespective of age groups (young + old). Dashed regression lines represent non-significant outputs of the GLMM.

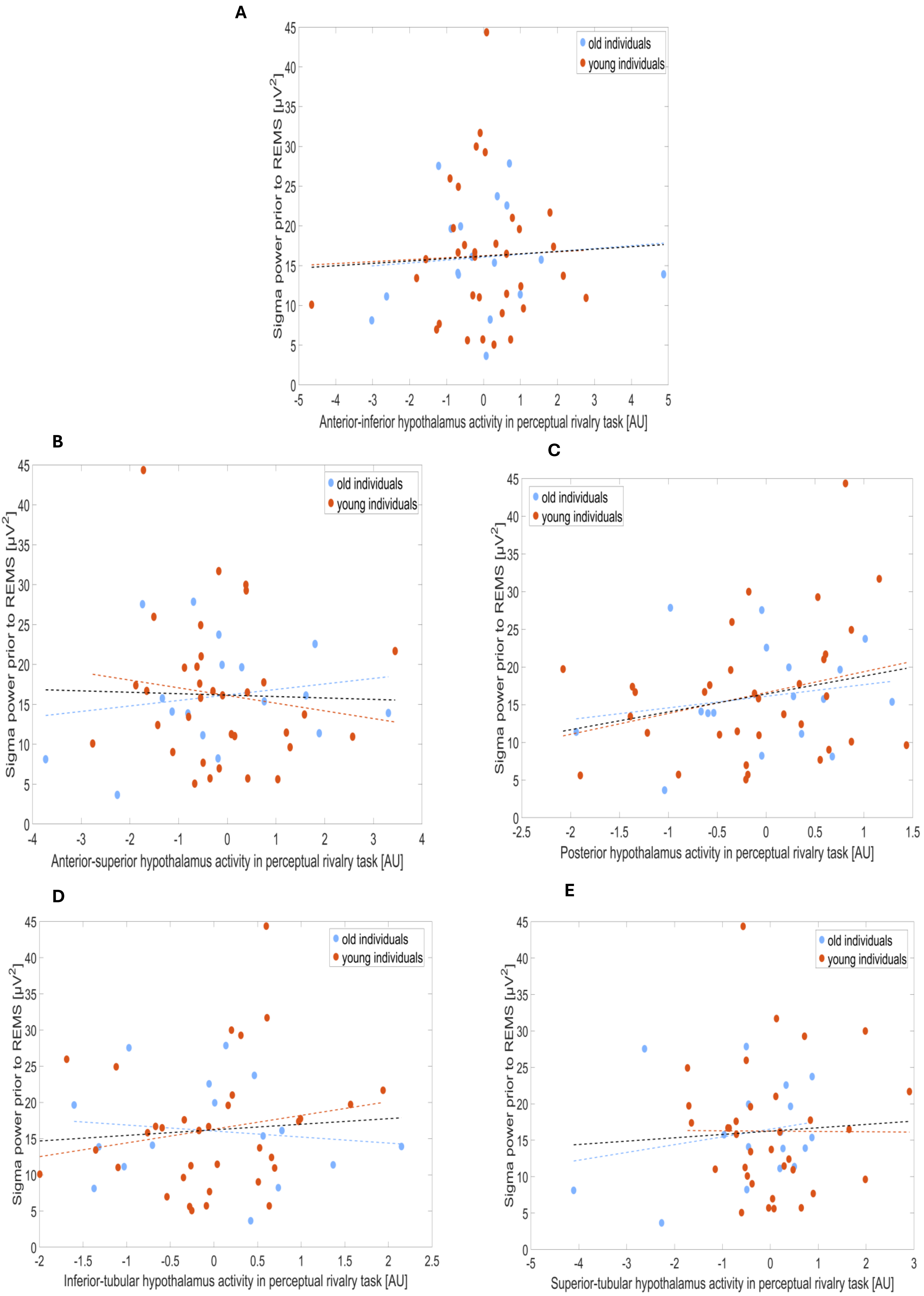
A
old individuals
young individuals
Sigma power prior to REMS [μV2]
Anterior-inferior hypothalamus activity in perceptual rivalry task [AU]
B
old individuals
young individuals
Sigma power prior to REMS [μV2]
Anterior-superior hypothalamus activity in perceptual rivalry task [AU]
C
old individuals
young individuals
Sigma power prior to REMS [μV2]
Posterior hypothalamus activity in perceptual rivalry task [AU]
D
old individuals
young individuals
Sigma power prior to REMS [μV2]
Inferior-tubular hypothalamus activity in perceptual rivalry task [AU]
E
old individuals
young individuals
Sigma power prior to REMS [μV2]
Superior-tubular hypothalamus activity in perceptual rivalry task [AU]

**Supplementary Figure S2. Non-significant associations between hypothalamus subparts activity estimates during the perceptual rivalry task and sigma power prior to REMS. (A)** Association between the interior-inferior hypothalamus activity estimates during the perceptual rivalry task and sigma power prior to REMS. **(B)** Association between the interior-superior hypothalamus activity estimates during the perceptual rivalry task and sigma power prior to REMS. **(C)** Association between the posterior hypothalamus activity estimates during the perceptual rivalry task and sigma power prior to REMS. **(D)** Association between the inferior-tubular hypothalamus activity estimates during the perceptual rivalry task and sigma power prior to REMS. **(E)** Association between the superior-tubular hypothalamus activity estimates during the perceptual rivalry task and sigma power prior to REMS.

The GLMM did not yield a statistical trend for the hypothalamus activity by hypothalamus subpart interaction (p=0.8).

Simple regression lines are used for a visual display and do not substitute the GLMM outputs. The black line represents the regression irrespective of age groups (young + old). Dashed regression lines represent non-significant outputs of the GLMM.

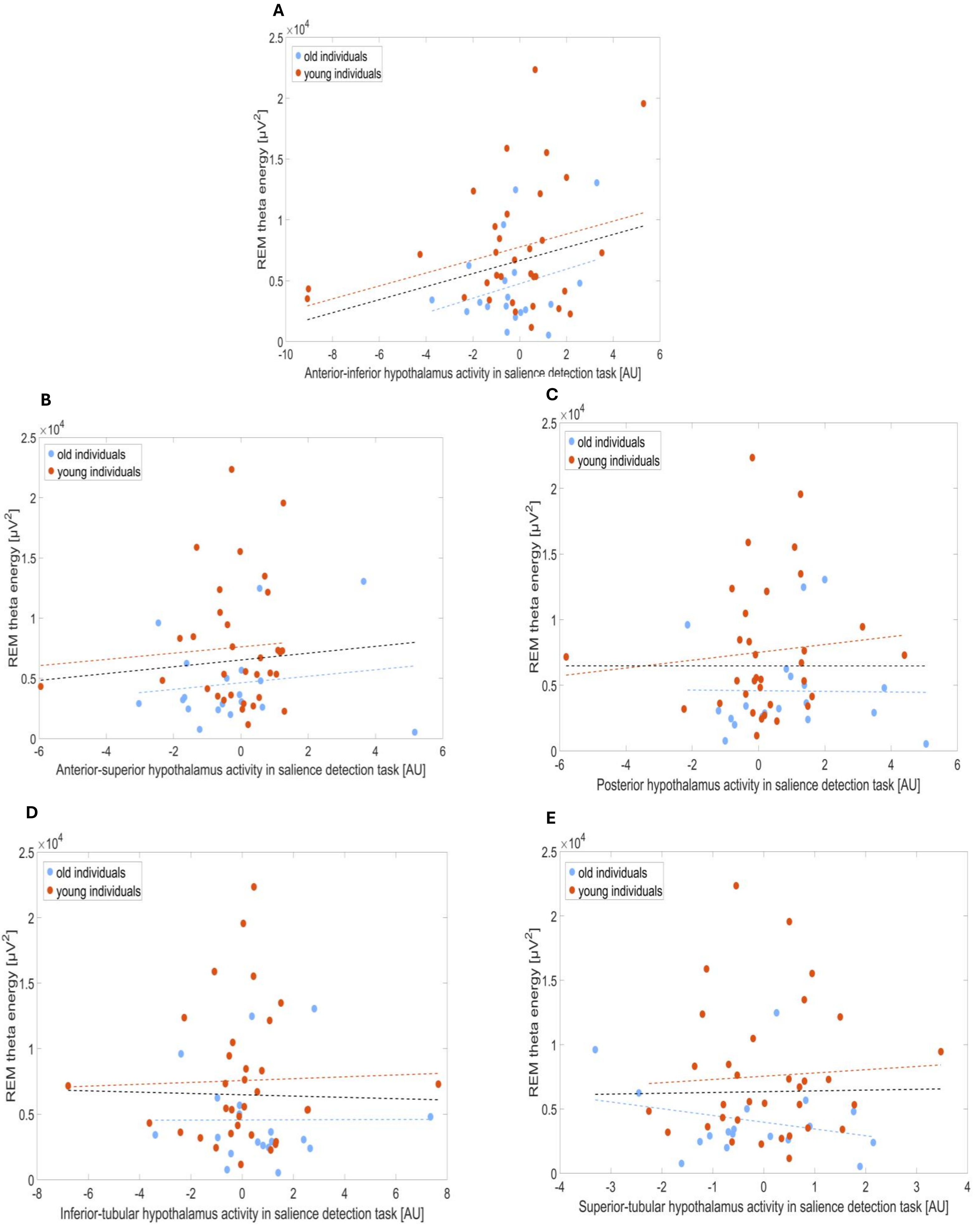
A
B
C
D
E
old individuals
young individuals
REM theta energy [μV$^2$]
$\times 10^4$
Anterior-inferior hypothalamus activity in salience detection task [AU]
Anterior-superior hypothalamus activity in salience detection task [AU]
Posterior hypothalamus activity in salience detection task [AU]
Inferior-tubular hypothalamus activity in salience detection task [AU]
Superior-tubular hypothalamus activity in salience detection task [AU]

**Supplementary Figure S3. Non-significant associations between hypothalamus subparts activity estimates during the salience detection task and REM theta energy. (A)** Association between the interior-inferior hypothalamus activity estimates during the salience detection task and REM theta energy. **(B)** Association between the interior-superior hypothalamus activity estimates during the salience detection task and REM theta energy. **(C)** Association between the posterior hypothalamus activity estimates during the salience detection task and REM theta energy. **(D)** Association between the inferior-tubular hypothalamus activity estimates during the salience detection task and REM theta energy. **(E)** Association between the superior-tubular hypothalamus activity estimates during the salience detection task and REM theta energy.

The GLMM did not yield a statistical trend for the hypothalamus activity by hypothalamus subpart interaction (p=0.9).

Simple regression lines are used for a visual display and do not substitute the GLMM outputs. The black line represents the regression irrespective of age groups (young + old). Dashed regression lines represent non-significant outputs of the GLMM.

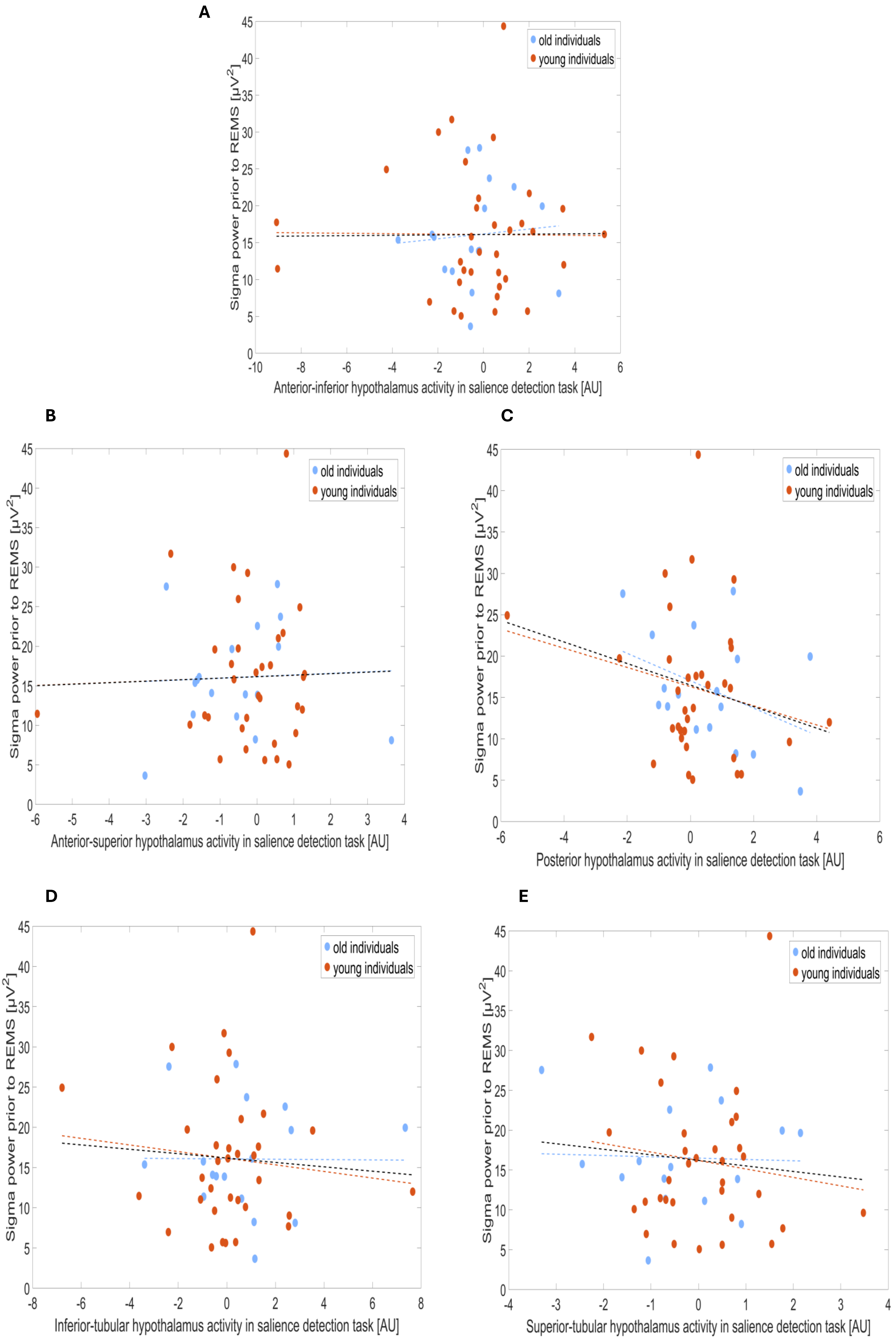
A
old individuals
young individuals
Sigma power prior to REMS [μV²]
Anterior-inferior hypothalamus activity in salience detection task [AU]
B
old individuals
young individuals
Sigma power prior to REMS [μV²]
Anterior-superior hypothalamus activity in salience detection task [AU]
C
old individuals
young individuals
Sigma power prior to REMS [μV²]
Posterior hypothalamus activity in salience detection task [AU]
D
old individuals
young individuals
Sigma power prior to REMS [μV²]
Inferior-tubular hypothalamus activity in salience detection task [AU]
E
old individuals
young individuals
Sigma power prior to REMS [μV²]
Superior-tubular hypothalamus activity in salience detection task [AU]

**Supplementary Figure S4. Non-significant associations between hypothalamus subparts activity estimates during the salience detection task and sigma power prior to REMS. (A)** Association between the interior-inferior hypothalamus activity estimates during the salience detection task and sigma power prior to REMS. **(B)** Association between the interior-superior hypothalamus activity estimates during the salience detection task and sigma power prior to REMS. **(C)** Association between the posterior hypothalamus activity estimates during the salience detection task and sigma power prior to REMS. **(D)** Association between the inferior-tubular hypothalamus activity estimates during the salience detection task and sigma power prior to REMS. **(E)** Association between the superior-tubular hypothalamus activity estimates during the salience detection task and sigma power prior to REMS.

The GLMM did not yield a statistical trend for the hypothalamus activity by hypothalamus subpart interaction (p=0.6).

Simple regression lines are used for a visual display and do not substitute the GLMM outputs. The black line represents the regression irrespective of age groups (young + old). Dashed regression lines represent non-significant outputs of the GLMM.

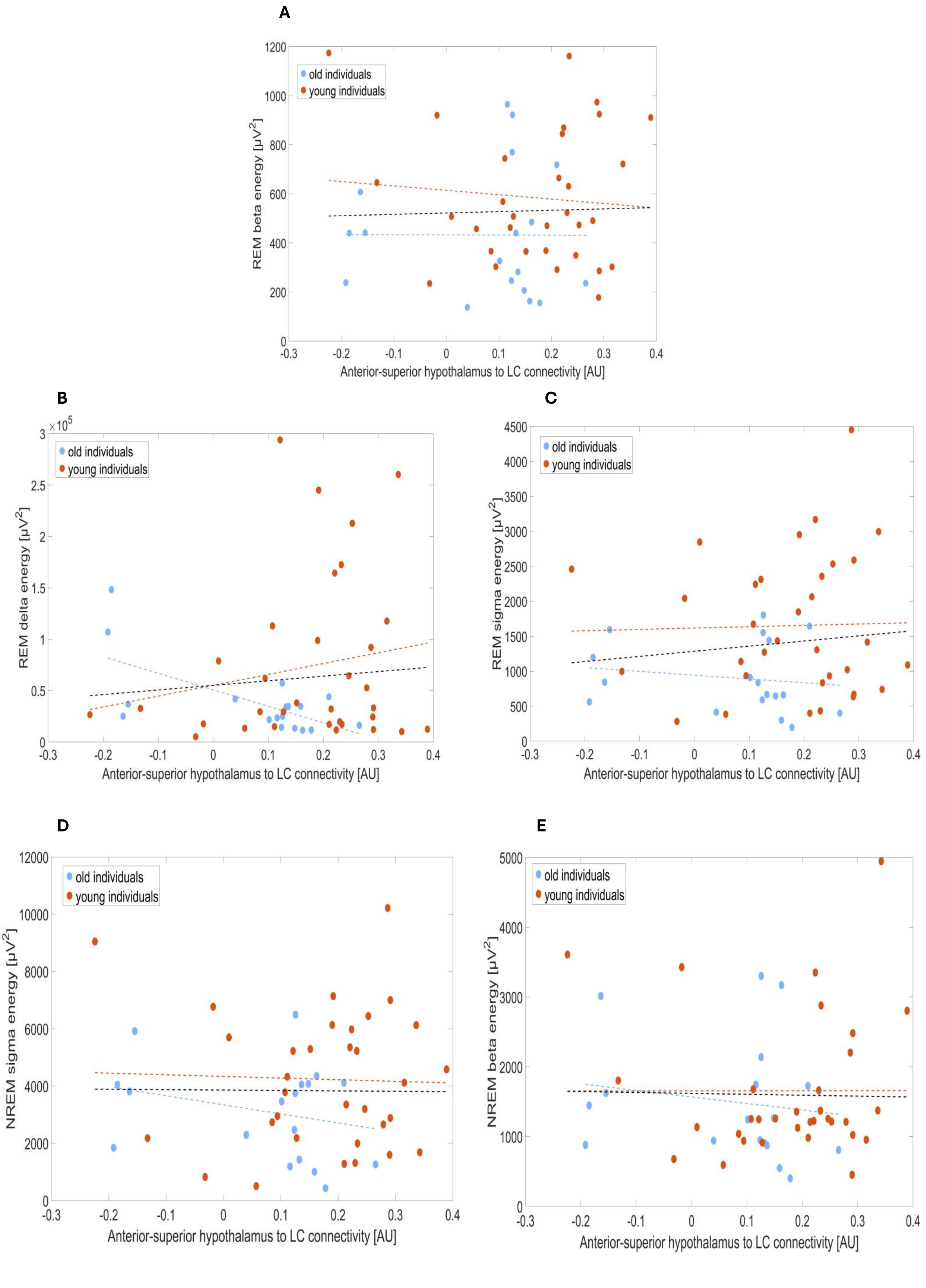
A
old individuals
young individuals
REM beta energy [μV²]
Anterior-superior hypothalamus to LC connectivity [AU]
B
REM delta energy [μV²]
C
REM sigma energy [μV²]
D
NREM sigma energy [μV²]
E
NREM beta energy [μV²]

**Supplementary Figure S5. Non-significant associations between sleep metrics of interest and the connectivity from anterior-superior hypothalamus to LC to test the specificity. (A)** Association between REM beta energy and the anterior-superior hypothalamus to LC connectivity. **(B)** Association between REM delta energy and the anterior-superior hypothalamus to LC connectivity. **(C)** Association between REM sigma energy and the anterior-superior hypothalamus to LC connectivity. **(D)** Association between NREM sigma energy and the anterior-superior hypothalamus to LC connectivity. **(E)** Association between NREM beta energy and the anterior-superior hypothalamus to LC connectivity.

None of the associations were significant ($p>0.051$).

Simple regression lines are used for a visual display and do not substitute the GLMM outputs. The black line represents the regression irrespective of age groups (young + old). Dashed regression lines represent non-significant outputs of the GLMM.
